# Laquinimod treatment attenuates EAU by inhibiting both the inductive and effector phases in an APC-dependent manner

**DOI:** 10.1101/2025.05.20.654165

**Authors:** Biying Xu, Mary J. Mattapallil, Xiuzhi Jia, Jihong Tang, Reiko Horai, Rachel R. Caspi, Igal Gery

## Abstract

**Purpose:** To evaluate the immunomodulatory effects on experimental autoimmune uveitis (EAU) of the aryl hydrocarbon receptor (AhR) agonist Laquinimod (LAQ), and its active metabolite DELAQ, with a focus on dendritic cell– and T cell–mediated mechanisms.

**Methods:** EAU was induced in mice by active immunization or by adoptive transfer of activated T cells. Mice were treated with LAQ either from the time of immunization or from 7 days after. Effects of LAQ were examined in wild-type, global AhR-knockout, or dendritic cell–conditional AhR knockout mice. Direct vs. indirect effects of AhR agonism on dendritic cells and T cells were studied *in vitro* using DELAQ, a major active metabolite of LAQ.

**Results:** LAQ treatment from Day 0 fully suppressed EAU, while delayed treatment (Day 7) provided only partial protection. In the adoptive transfer model, LAQ-treated recipients showed reduced pathology. Global AhR-deficient mice developed severe EAU comparable to that of wild-type mice, with an elevated Th17 response. LAQ-treated mice displayed increased frequencies of cDC1 and FoxP3⁺ regulatory T cells. In vitro, DELAQ activated AhR signaling and induced *Ido1* and *Ido2* expression in dendritic cells. DELAQ inhibited the activation of naïve and memory mouse T cells in an APC-dependent manner, as the response to anti-CD3/CD28 stimulation was unaffected. Importantly, DELAQ suppressed recall responses of human PBMC to tetanus toxoid.

**Conclusions:** LAQ protects against EAU by acting on AhR-expressing antigen-presenting cells to impair both priming and reactivation of pathogenic T cells. Its active metabolite DELAQ does not suppress T cells directly, but re-programs dendritic cells toward a tolerogenic, IDO-expressing phenotype that promotes immune regulation.

## Introduction

Noninfectious uveitis is an autoimmune inflammation of the retina and uvea caused by activated autoreactive T cells specific to retinal antigens, and is amplified by recruited leukocytes^1,2^. Cellular autoimmune processes play an essential role in the pathogenesis of the disease. Experimental autoimmune uveitis (EAU), a pre-clinical model of uveitis, is induced by T cells specific for retinal antigens^3–5^. In mice, EAU can be induced by immunization with IRBP or its immunogenic peptides. Alternatively, it can be generated by the adoptive transfer of activated T cells, isolated from immunized mice, into naïve recipient mice^6^.

The aryl hydrocarbon receptor (AhR) is a ligand-activated transcription factor. Its activation regulates complex biological processes and maintains the skin, lungs, and gut barrier functions^7^. Its expression in various immune cells contributes to immune function. AhR ligands, including dietary and environmental stimulants, microbiome metabolites, and endogenous molecules, exhibit ligand-specific, cell-type-specific, and species-specific transcriptional regulation^8,9^.

Myeloid cells and regulatory T cells express high levels of AhR and represent a target population for AhR ligands.^10^ Dendritic cells (DCs), a critical component of professional antigen-presenting cells (APCs), play a central role in shaping T-cell responses and immune tolerance^11^. Interestingly, AhR activation influences both the differentiation and function of DCs, potentially explaining its immunomodulatory effect. AhR activation in DCs can cause an immunosuppressive effect by inducing the expression of indoleamine 2,3-dioxygenase (IDO) 1 and 2 enzymes. IDO1 and IDO2 catalyze kynurenine production from tryptophan metabolism, which promotes Foxp3+ Treg differentiation^12^. The IDO1-Kyn-AhR axis, a metabolic signaling pathway, can also facilitate a tolerogenic phenotype among conventional DC subsets^13,14^.

Laquinimod (LAQ) is a pro-drug whose active metabolite, DELAQ, functions as an AhR agonist. It has previously been demonstrated to suppress experimental autoimmune encephalomyelitis (EAE), a model for multiple sclerosis^15–17^, which shares essential immunological mechanisms with EAU. Furthermore, LAQ has been assessed in several clinical trials for multiple sclerosis ^18^ and Huntington’s disease^19^, with limited or inconclusive success. Although the exact mechanism of action is not fully understood, several mechanisms have been proposed^20^, including reprogramming APCs to shift the balance between T effectors and regulatory T cells^21–23^, tightening the blood-brain barrier by inducing junction protein expression, which reduces pathogenic cell invasion^24^, and modulating brain-derived neurotrophic factor (BDNF) to provide neuroprotective activity^22,25^ . *In vitro* experiments using DELAQ demonstrated activation of the same AhR-driven molecular pathways associated with the protective effects of LAQ observed *in vivo*^17^.

In the current study, we examined the effects of LAQ treatment on induction of EAU and on modulating its associated immune processes. To dissect the effect of DELAQ on immune cells in vitro, we focused on dendritic cells and T cells, whose function is central in the pathogenesis of EAU development. Data using AhR conditional knockout mice supported the conclusion that LAQ’s protective effect in EAU is primarily mediated via modulation of dendritic cells, with what could be indirect effects on T-cell function.

## Materials and Methods

### Mice

C57BL/6J (B6), OTII, AhR^fl/fl^, and CD11c-Cre mice were purchased from Jackson Lab (Bar Harbor, ME). AhR-null mice were provided by Dr. Gonzalez (NCI, NIH)^26^. Mice were housed in a specific pathogen-free facility with access to standard facility chow and water *ad libitum*. All procedures were conducted in compliance with ARVO policy and NIH guidelines on the care and use of animals in research. All study protocols were approved by the Animal Care and Use Committee (ACUC) of the National Eye Institute.

### EAU induction and evaluation

B6 mice were immunized by subcutaneous injection with 300 µg of human interphotoreceptor retinoid-binding protein (IRBP) peptide 651-670 (IRBP_651-670_, emulsified in complete Freund’s adjuvant).^27^ Pertussis toxin, one µg (Sigma, St Louis, MO), was injected intraperitoneally. LAQ, provided by Active Biotech AB (Lund, Sweden), was dissolved in PBS and administered daily by oral gavage at 25 mg/kg; control mice received daily gavage of PBS. Two different treatment regimens were tested; mice were either treated from the day of immunization (Day 0 treatment) or day 7 post-immunization (Day 7 treatment) until day 21. Fundoscopy was used to monitor EAU development from day 9 to day 21. Disease scores ranging from 0 to 4 were assigned to each eye^3^, and the average scores of the two eyes of each mouse were used to determine EAU progression.

Mice were euthanized on day 21 post-immunization. Their eyes were collected, fixed, dehydrated, and embedded in methacrylate or paraffin before being sectioned through the optic nerve plane. This was followed by hematoxylin and eosin staining for histological analysis. Pathological changes were evaluated using a scale of 0 to 4^4^.

For the adoptive transfer of EAU, activated IRBP_651-670_ specific T cells were transferred to naïve B6 or congenic CD45.1 mice.^12^ WT C57BL/6J or AhR-null mice on C57BL/6J background were immunized with IRBP_651-670_ emulsified in CFA and one µg of pertussis toxin. Donor mice were euthanized on day 14. Splenocytes, inguinal lymph node cells (which drains the site of immunization), and eyes were collected and were cultured with IRBP_651-670_ for 72 hours before being transferred to recipient mice. Fifty million cells were transferred to each recipient via intraperitoneal injection. The recipient mice received daily oral gavage of LAQ at 25 mg/kg or PBS for control, starting the day before adoptive cell transfer and continuing until day 11. The disease progression was monitored by fundoscopy from day 5 until day 11 post-cell transfer. Mice were euthanized on day 11, and their eyes were collected and analyzed for histopathological changes.

### Analysis of cellular immune responses

*Lymphocyte recall proliferative response*. Active immunized mice were euthanized on day 21 post-immunization. Splenocytes were cultured with IRBP_651-670_, and the stimulation level was determined by a conventional [3H]-thymidine incorporation assay^27^. Some cultures were treated with DELAQ (provided by Active Biotech) at a final concentration of 100 ng/ml.

*Cytokine ELISAs.* Culture supernatants were collected from splenocyte culture after 48 hours, except for the supernatant for IL-10 ELISA, which was collected after 72 hours. The levels of secreted cytokines were determined using commercial ELISA kits from R&D Systems (Minneapolis, MN): mouse IL-2 DuoSet ELISA kit (Cat. Nr DY402); mouse IL-4 DuoSet ELISA kit (Cat. Nr DY404); mouse IL-6 DuoSet ELISA kit (Cat. Nr DY406); mouse IL-17 DuoSet ELISA kit (Cat. Nr DY421); mouse IL-22 DuoSet ELISA kit (Cat. Nr DY582); mouse TNF-alpha DuoSet ELISA kit (Cat. Nr DY410); mouse IFN-gamma DuoSet ELISA kit (Cat. Nr DY-485); and mouse IL-10 Quantikine ELISA kit (Cat. Nr S1000B).

*FACS analysis.* FACS analysis was used to assess intracellular cytokine and transcription factor expression. Splenocytes and cells from the draining inguinal lymph node (DLN) were used for intracellular staining of cytokines IFN-γ, IL-17, and transcription factors AhR, Foxp3, T-bet, and RORγt.

For intracellular cytokine staining, cells were stimulated with PMA (50ng/ml), ionomycin (500ng/ml), and Brefeldin A (GolgiPlug, BD) for 4 hours. This was followed by 4% paraformaldehyde fixation and BD permeabilization buffer (BD Cat. No. 51-2091KZ) before staining for IFN-γ and IL-17A overnight. For staining of transcription factors (AhR, T-bet, RORγt and Foxp3) the eBioscience Foxp3/transcription factor staining buffer set (Cat. No. 00-5523-00) from Invitrogen was used according to the manufacturer’s protocol.

To characterize conventional DC populations, spleens were digested with collagenase at 37°C for 30 min before surface staining for 30 minutes at 4°C, to identify cDC1 and cDC2.

Briefly, the CD11c+ MHCII+ DC population was gated by initially excluding CD19+ MHCII+ B cells and subsequently excluding F4/80+ CD64+ macrophage populations. Among CD11c+ MHCII+ cells, XCR-1+ cells were identified as cDC1, while CD11b+ cells were identified as cDC2. Cells were incubated using Fc block (2.4G2, BioXcell) and stained with the following commercial antibodies: anti-CD3 (clone 145-2C11), anti-CD4 (RM4.5), anti-CD8 (53-6.7) anti-CD25 (PC61.5), anti-CD45 (30-F11), IFN-γ (XMG1.2), IL-17A (TC11-18H10.1), AhR (4MEJJ), T-bet (4B10), RORγt (AFKJS-9), Foxp3 (FJK-16S), Anti-MHCII (M5/114.15.2), anti-CD19 (1D3), anti-F4/80 (BM8), anti-CD64 (X54-5/7.1), anti-CD11c (N418), anti-CD11b (M1/70), and anti-XCR1(ZET). Samples were analyzed using a Cytoflex flow cytometer (Miltenyi) and FlowJo software (Tree Star, Ashland, OR).

### Quantitative PCR

Cells and tissue samples were lysed in Trizol reagent (Sigma Aldrich). RNA was purified using a RNeasy mini kit (Qiagen). A high-capacity cDNA Reverse Transcription kit (Thermo Fisher Scientific) was utilized for cDNA synthesis. TaqMan real-time PCR Assay and TaqMan Fast Universal PCR Master Mix (Thermo Fisher Scientific) were used. Data were acquired using the Quant Studio 7 Flex q-PCR machine (Applied Biosystems) and analyzed using the ΔΔCt method for relative quantification (RQ) with *Actb* as the endogenous control and untreated cells as the reference sample.

TaqMan real-time PCR Assays used were as follows: *Actb*: Mm02619580_g1 *Ahrr*: Mm00477443_m1 *Cyp1a1*: Mm00487218_m1 *Cyp1b1*: Mm00487229_m1 *Ido1*: Mm00492586_m1 *Ido2*: Mm00524206_m1

### Bone Marrow-derived Dendritic Cells (BMDCs) generation

Bone marrow cells were flushed from the femur and tibiae of B6 or AhR-null mice following the standard protocol^28^. Red blood cells were lysed using ACK lysing buffer. For dendritic cell (DC) differentiation, bone marrow cells were cultured in 100 mm Petri dishes at a density of 1 x 10^^6^ cells per ml in RPMI, supplemented with 10% FBS (Hyclone) and 20 ng/ml of GM-CSF (315-03, Peprotech). The culture medium was replaced on days 3, 5, and 8. BMDCs were collected on day 9 for activation with LAQ or DELAQ at a final concentration of 100 ng/ml for 24 hours.

### Naïve T cell activation and polarization

Naïve CD4+ T cells were isolated from the splenocytes of B6 or AhR-null mice using the EasySep mouse naïve CD4+ T cell isolation kit (StemCell Technologies). The enriched CD4+ T cells were activated for 48 hours with plate-bound anti-CD3 (1µg/ml, clone 145-2C11, BioXcell) and anti-CD28 (0.5µg/ml, clone 37.51, BioXcell) antibodies, and cultured in RPMI supplemented with 10% fetal bovine serum (FBS). We tested LAQ or DELAQ at 100 ng/ml. A conventional [^3^H]-thymidine incorporation assay was used to determine the T cell proliferative response.

We used combinations of various cytokines and neutralizing antibodies for the T-cell polarization assay. The polarization conditions are as follows: Th17: anti-IFN-γ (10 μg/ml; R4-6A2), anti-IL-4 (10 μg/ml; 11B11), TGF-β (2.5 ng/ml), IL-6 (25 ng/ml), and IL-23 (10 ng/ml); Th1: anti-IL-4 (10 μg/ml; 11B11) and IL-12 (10 ng/ml); Treg: TGF-β1 (5 ng/ml) and IL-2 (10 ng/ml). Cells were stimulated in different polarization conditions with or without DELAQ for 72 hours before being analyzed by flow cytometry.

### Mouse T-cell priming assay

Irradiated (30 Gy) splenocytes from B6 mice were plated at 5 × 10^5 cells per well in 96-well plates. Naïve CD4+ T cells from OT-II mice were purified using the EasySep Mouse Naïve CD4+ Isolation Kit (Stem Cell Technologies) and plated at 1 × 10^5 cells per well. These cells were stimulated with OVA peptide 323–339 at 10 µg/ml. 100 ng/ml of DELAQ was added to the culture mix. After 48 hours of incubation, the plates were pulsed with ^3^H thymidine and harvested 16 hours later.

### Recall response to Tetanus Toxoid of human PBMCs

PBMCs were isolated from the buffy coats of healthy adult donors obtained from the NIH blood bank. PBMCs were recovered by centrifugation using Accuspin Histopaq-1077 (Sigma Cat# A7054). The cells were washed twice with PBS and resuspended in RPMI medium supplemented with 5% human AB serum. PBMCs (5 × 10^6/ml) were stimulated with Tetanus Toxoid (List Biologics) at 1 µg/ml, and the effect of DELAQ on cell proliferation was tested at 100 ng/ml.

## Results

### LAQ treatment attenuates EAU development

Mice that were actively immunized to induce EAU and received LAQ treatment from Day 0, were fully protected against disease induction; those treated from Day 7 exhibited lower disease scores (Fig. 1A). In agreement with the fundoscopy scores, histological analysis revealed moderate to severe disease in mice treated with PBS, regardless of the treatment regimen. Mice that underwent LAQ treatment from Day 0 displayed minimal retinal damage and immune cell infiltration (Fig. 1B). Reduced, but still statistically significant protection was noted in mice treated from Day 7 (Fig. 1C). We conclude that LAQ treatment is effective in inhibiting the development of EAU in this model.

**Figure 1.**
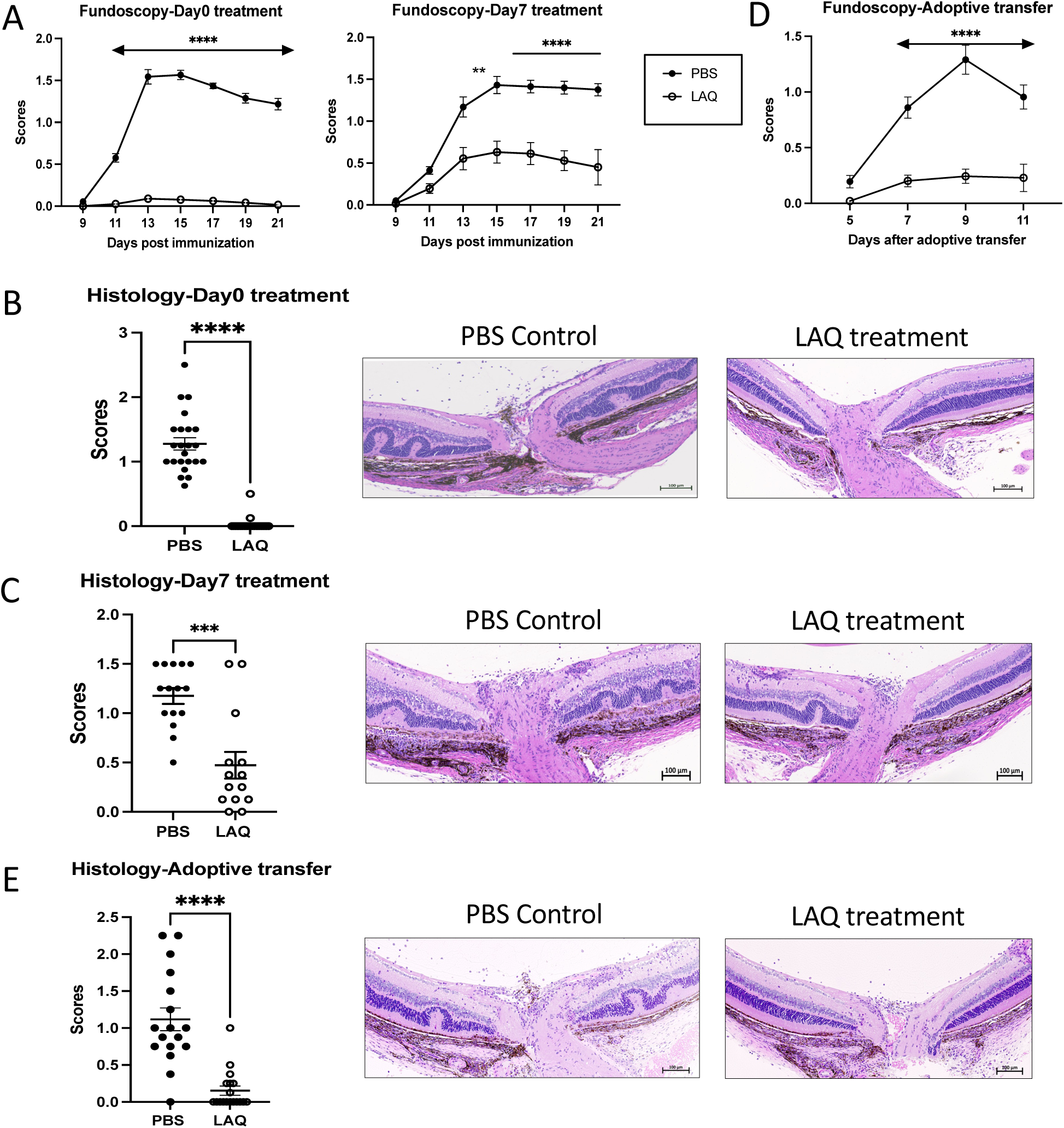
LAQ treatment attenuates EAU in the inductive and effector phases LAQ was administered by gavage to actively immunized or adoptively transferred EAU mice daily at 25mg/kg. For actively immunized EAU, mice received LAQ treatment from Day 0 or Day 7 post-immunization. For adoptively transferred EAU, mice received LAQ treatment from the day before cell transfer. **A -** Fundoscopy scores of immunized mice were evaluated every two days from day 9 postimmunization. Fundoscopy scores of Day 0 treatment (combined data of four experiments) and Day 7 treatment group (combined data from four experiments). **B and C -** Histology scores of eyes collected on day 21 post-immunization. Representative histopathology images of the uveitic retina with H&E stain. **D -** Fundoscopy scores of adoptively transferred EAU mice were evaluated starting from day 5 post-cell-transfer (combined data of four experiments). **E -** Histology scores of eyes collected on day 11 post-cell-transfer, with representative histopathology images of the uveitic retina with H&E stain. All data were plotted as mean ± SEM, and Two-way ANOVA (A, D) and Mann-Whitney test (B, C, E) were used to determine the significance. *p <0.05, **p <0.01, ***p <0.001.

We next assessed the effect of LAQ on already activated effector T cells by using the model of adoptively transferred EAU. Activated T cells caused moderate to severe disease in PBS-treated recipients which was strongly inhibited by LAQ-treatment (Fig. 1D, E). Together, these findings suggest that LAQ treatment protects from EAU by inhibiting the development as well as the effector function of pathogenic T cells *in vivo* and attenuates EAU in both inductive and effector phases.

### Th1 and Th17 responses are inhibited by LAQ treatment, and are associated with an increase in Foxp3+ regulatory T cells

The molecular target of LAQ is AhR^16^. As proof of principle, we therefore tested AhR expression in T cells of LAQ-treated mice by flow cytometry. A decreased proportion of AhR-expressing CD4+ T cells was detected in the spleen and draining lymph nodes of mice treated with LAQ from day 0 (Fig. S1A). Since activated pathogenic T cells express AhR, the decreased percentage of AhR-expressing cells after LAQ treatment suggests an effect on the generation of pathogenic T cells *in vivo*.

To evaluate the effect of LAQ on the generation of Th1 and Th17 effector cells, we examined the antigen-specific responses of lymphocytes from immunized and LAQ-treated mice. We observed a reduced proliferative response to IRBP_651-670_ in LAQ-treated mice. The inhibitory effect was more pronounced in mice treated from day 0 than in those treated from day 7 (Fig. 2A). The lymphocyte culture supernatants from LAQ-treated mice showed a reduction in both IFN-γ and IL-17 (Fig. 2B). Consistent with the reduced antigen-specific IFN-γ and IL-17 cytokine production, flow cytometry analysis revealed a decrease in both Th1 and Th17 cell populations in the spleen and inguinal lymph nodes of LAQ-treated mice (Fig. 2C). Thus, the protective effect of LAQ may be due to impaired antigen-specific Th1 and Th17 effector T cell function.

**Figure 2.**
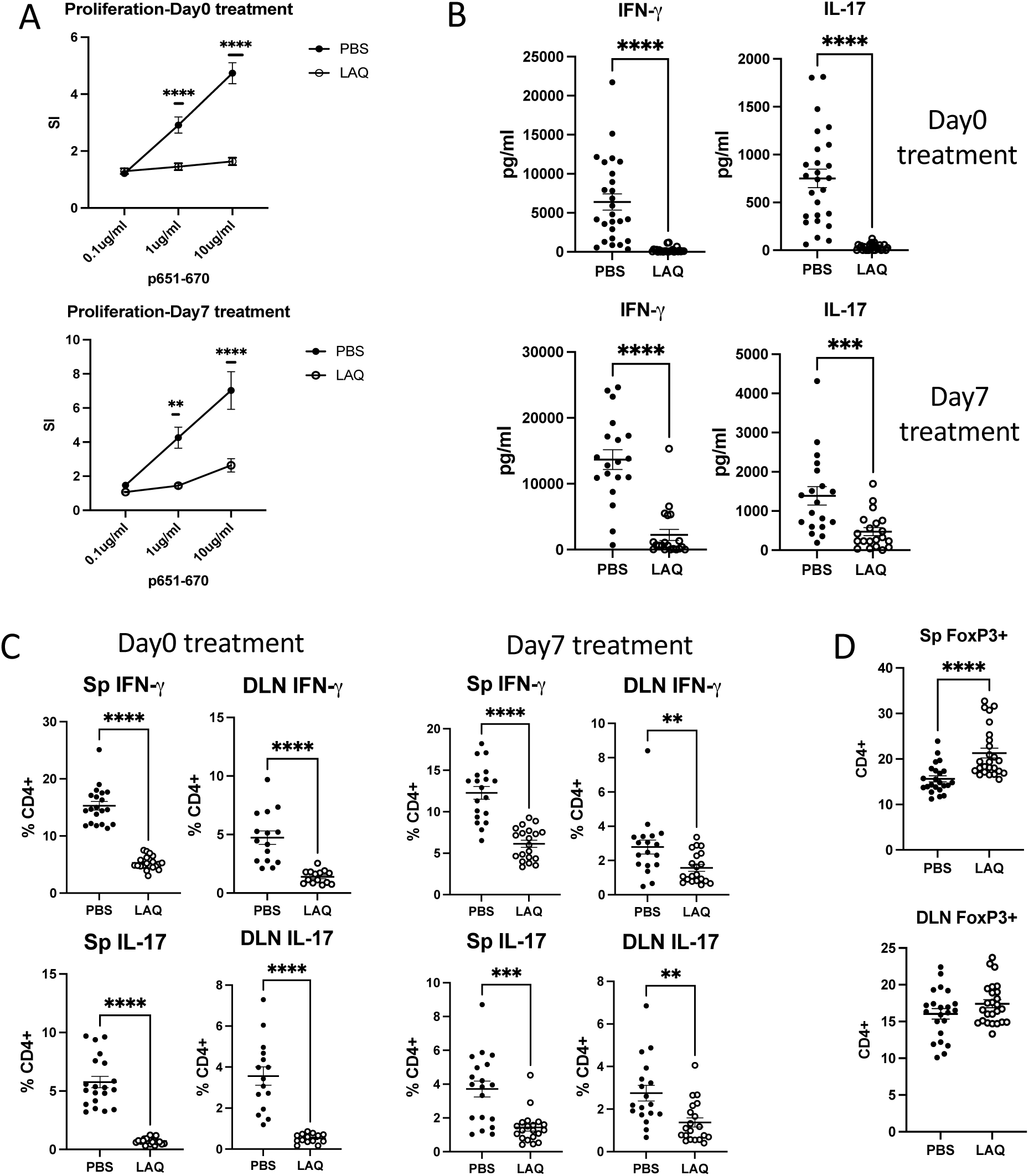
LAQ treatment inhibits Th1/Th17 responses and increases FoxP3⁺ regulatory T cells Mice with EAU were euthanized on day 21 after immunization. Cells from the spleen (SP) and draining lymph nodes (DLN) were purified for *ex vivo* testing. **A -** Antigen-specific (IRBP p651-670) proliferative response of splenocytes was plotted as stimulation indices (SI) - combined data of four experiments from day 0 or day 7 treatment groups. **B -** IFN-γ and IL-17 ELISA on culture supernatants of splenocytes stimulated with p651-670. **C -** Intracellular IL-17A and IFN-γ were detected by flow cytometry following *ex vivo* 4 h stimulation with PMA and ionomycin in the presence of Brefeldin. Frequency of IFN-γ, IL-17A-producing cells in total CD4+ T cells in the spleen, DLN of day0 and day7 treated groups. **D -** Intracellular Foxp3 was tested by flow cytometry without stimulation. Each dot represents one mouse, and all data were plotted as mean ± SEM. Mann-Whitney test was used to determine the significance. *p <0.05, ** p <0.01, *** p <0.001, ****p <0.0001.

We observed a reduction in pro-inflammatory cytokines (TNF-α and IL-6) and the Th2 cytokine (IL-4) in the splenocyte culture supernatants of day 0 LAQ-treated mice (Fig. S1B).

Interestingly, the antigen-induced production of IL-2, a critical cytokine for immune activation, and anti-inflammatory cytokines such as IL-10 and IL-22 were also lower in LAQ-treated mice (Fig. S1B). Although a reduction in IL-10 appears inconsistent with the increased Treg population, it could reflect a regulatory mechanism independent of IL-10 production.

An increased population of Foxp3+ regulatory T cells was observed in splenocytes but not in inguinal lymph node cells collected on day 21 in mice treated with LAQ from day 0 (Fig. 2D). It has been suggested that Foxp3+ regulatory T cells migrate from lymph nodes to target tissues to exert their regulatory function. Therefore, it is unsurprising that a change in the Treg population was not observed in draining lymph nodes at the end of the disease course.

Furthermore, an increased Treg population in the spleen was detected only in mice treated from day 0 (Fig. 2D) and not in those treated from day 7 (data not shown).

### Global deficiency of AhR permits full development of EAU

To understand how AhR activation influences EAU development, we examined the disease course in AhR-deficient mice (AhR-null mice). Immunized AhR-null mice developed EAU with scores similar to B6 mice (Fig. 3A). Histological analysis of pathological retinal changes revealed no differences between AhR-null mice treated with LAQ or PBS (Fig. 3B). These observations suggest that global deficiency of AhR does not significantly affect EAU development, which is in accordance with published data in EAE^16^. In line with this, antigen-specific proliferative responses to IRBP_651-670_ did not differ between LAQ-treated AhR deficient mice and PBS-treated WT or AhR-null mice (Fig. 3C). In contrast, however, AhR-null mice had increased IL-17 and decreased IL-10 levels compared to WT mice, which were unaffected by LAQ treatment, consistent with the AhR dependency of these responses. On the other hand, IFN-γ was minimally affected (Fig. 3D).

**Figure 3.**
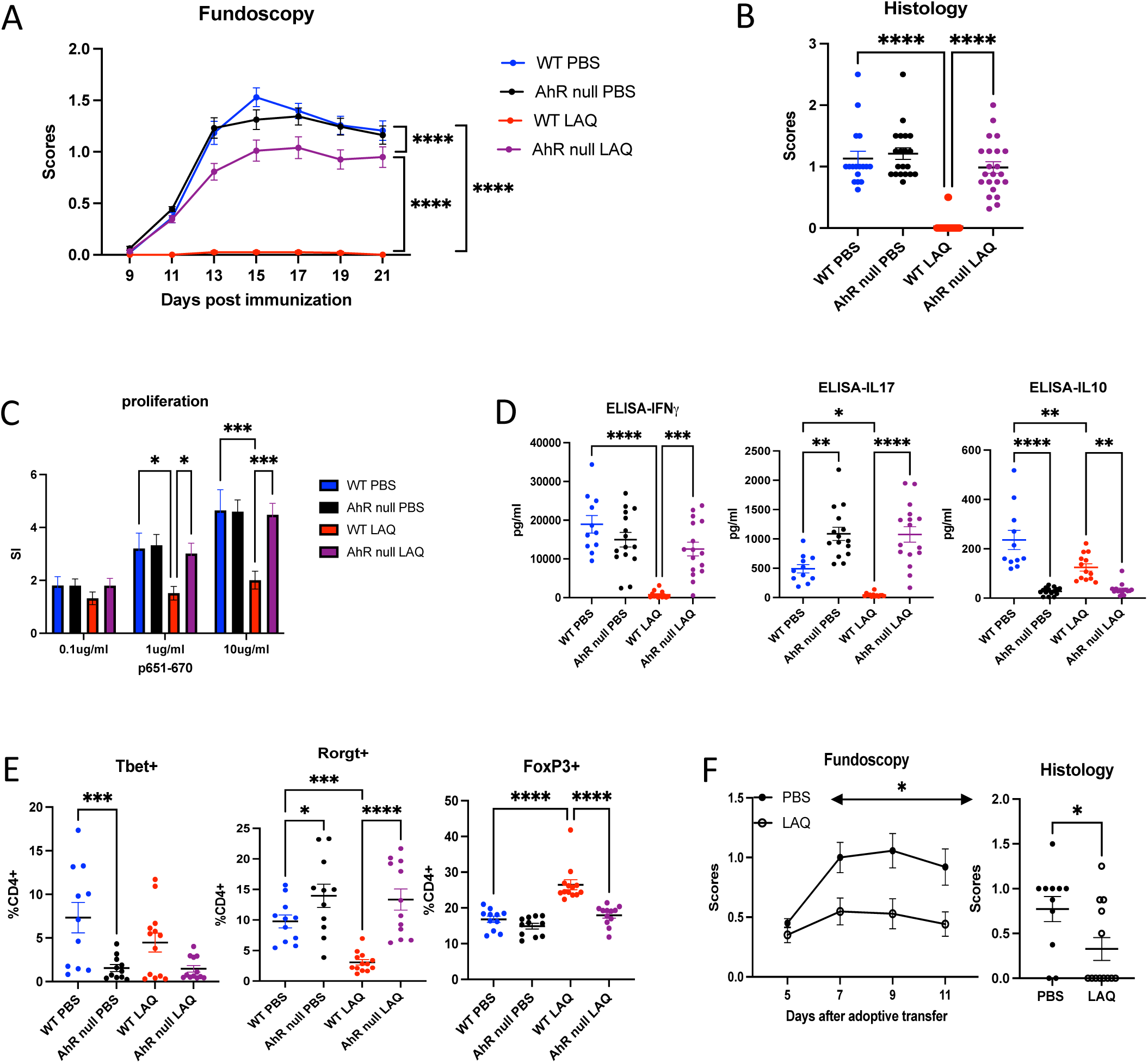
Global deficiency of AhR permits full development of EAU Immunized WT or AhR null mice were treated with LAQ from day 0. EAU development was evaluated by fundoscopy starting from day 9 till day 21 post-immunization. **A -** Fundoscopy scores. **B -** Histology scores. **C -** Antigen-specific (IRBP p651-670) proliferative response of splenocytes was plotted as stimulation indices (SI). **D -** Culture supernatants from splenocytes stimulated with p651-670 were tested for IFN-γ, IL-17, and IL-10 by ELISA. **E -** Lymphocytes were isolated from the spleen (SP) and analyzed by flow cytometry for T-bet, RORγt, and Foxp3 expression. **F -** Fundoscopy and Histology evaluation of adoptive transfer EAU in CD45.2 WT recipients.

To better understand whether the altered IL-17 and IL-10 were regulated at the transcription level, we tested T-bet, RORγt, and Foxp3 expression in splenocytes and cells from inguinal lymph nodes. In agreement with the increased IL-17 detected by ELISA, splenocytes expressed higher levels of RORγt (Fig. 3E) and the same pattern was observed in cells collected from the inguinal lymph nodes (data not shown).

Consistent with their uveitogenic effector activity in the original donor, AHR deficient T cells of AhR-null donor mice induced moderate to severe adoptively transferred EAU in congenic CD45.1 recipient mice. These observations confirm that T cells from AhR-null mice can be primed and are pathogenic for EAU induction. Interestingly, LAQ-treated recipients of AHR-null effector T cells mice exhibited protection from EAU, as noted by fundoscopy and supported by histological analysis (Fig. 3F).Since the donor T cells originated from AhR-null donor mice, there should be no direct LAQ effect on the transferred T cells, and the protective effect of LAQ must have been due to effects on the host cells.

### DELAQ stimulation activates genes related to the AhR pathway and has differential effects in T cells and dendritic cells

As a pro-drug, LAQ exerts its effects *in vivo* after being metabolized by the liver, but is minimally effective *in vitro*. One of the known active metabolites of LAQ, DELAQ, was tested on T cells and dendritic cells (DCs) *in vitro*. DELAQ stimulated naïve CD4+ T cells showed elevated expression of AhR-pathway-related genes: *Ahrr*, *Cyp1a1*, and *Cyp1b1*, but not in cells from AhR-null mice (Fig. S2A), confirming that AhR is the molecular target of DELAQ. Of note, DELAQ did not affect T cell proliferation in response to CD3/CD28 stimulation (Fig. S2B), suggesting that their effect on T cell responses to, is indirect.

Next, we tested various T-cell polarization conditions (Th17, Th1, Treg) in the presence of DELAQ. Under Th17-polarizing conditions, DELAQ treatment enhanced Th17 differentiation (Fig. S2C). It is well-documented that AhR plays a crucial role in Th17 differentiation^29^; therefore, as an AhR ligand, it is not surprising that DELAQ promotes Th17. Similarly, DELAQ stimulation induced the AhR pathway-related genes *Cyp1a1* and *Cyp1b1* in BMDC (Fig. S2D). IDOs are enzymes essential for the Kynurenine pathway, which induce regulatory T cells and inhibit T effector functions. The expression of the *Ido1* and *Ido2* genes, well-known molecules for inducing tolerogenic DC, increased in BMDC from WT mice treated with DELAQ (Fig. S2E), in immature and mature DCs induced by TNF-α. Interestingly, *Ido* gene expression was also dependent on AhR upon *in vitro* DELAQ stimulation. These results support a scenario in which LAQ treatment *in vivo* upregulates the IDO/Kynurenine pathway in APCs to inhibit effector T cell function. Notably, we observed an increased expression of AhR-pathway-related genes upon DC maturation. AhR stimulation by DELAQ may have a greater effect on tolerogenic DC generation in the mature DC population.

Next, we generated a mouse strain in which AhR is conditionally knocked out specifically in the CD11c+ population, primarily composed of dendritic cells, the main players among professional APCs (Fig. 4A). Mice with AhR conditional knockout in the dendritic cell population (CD11cCre-AhR^fl/fl^) develop full-blown EAU that is still inhibitable by LAQ (Fig. 4B). CD11c AhR⁻ and their CD11c AhR⁺ littermates developed comparable EAU upon immunization. CD11c AhR⁻ mice treated with LAQ were partially protected from EAU induction compared to CD11c AhR⁺ mice (Fig. 4B). We detected the same trend in antigen-specific proliferative response and cytokine production in CD11c AhR⁻ mice treated with LAQ compared to CD11c AhR+ mice counterparts (Fig. 4C, 4D). Furthermore, LAQ treatment of EAU mice resulted in a reduction of the IL-17-promoting transcription factor RORγt in splenic CD4+ T cells, irrespective of AhR expression in CD11c+ DCs, albeit with varying degrees of suppression (Fig. 4E). In contrast, the expression of the IFNγ-promoting transcription factor T-bet in these cells remained unchanged.

**Figure 4.**
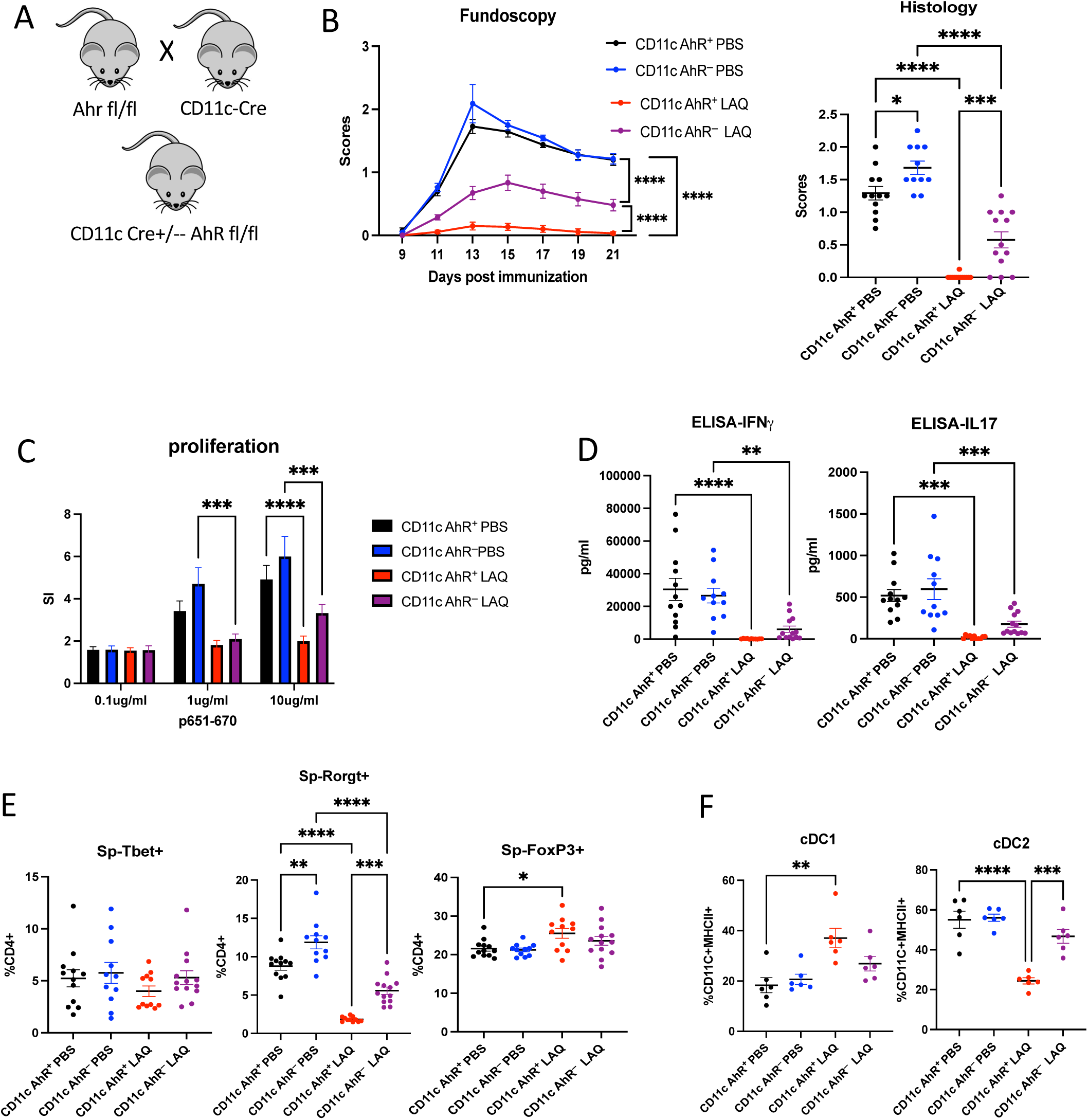
LAQ treatment partially protects CD11c-AhR-deficient mice from EAU Immunized CD11c AhR+ and CD11c AhR-littermates were treated with LAQ from Day 0. EAU development was evaluated by fundoscopy; mice were euthanized on day 21 post-immunization. **A -** Scheme on CD11c-cre AhR^fl/fl^ mouse strain generation. **B -** Fundoscopy and histology scores. **C -** The IRBP p651-670-specific proliferative response of splenocytes was plotted as stimulation indices (SI). **D -** IFN-γ, and IL-17 ELISA data on culture supernatants from splenocytes stimulated with IRBP p651-670. **E -** Splenocytes were isolated and analyzed by flow cytometry for T-bet, RORγt, and Foxp3 expression. **F -** Flow cytometry analysis on splenocytes for cDC1 and cDC2 populations. Each dot represents one mouse, and all data were plotted as mean ± SEM. Two-way ANOVA and Mann-Whitney test were used to determine the significance. *p <0.05, **p <0.01, ***p <0.001, ****p <0.0001

Absence of AhR in CD11c⁺ cells did not affect the severity of EAU in PBS-treated mice, indicating that AhR signaling in dendritic cells is not essential for disease induction. However, LAQ-treated CD11c-AhR⁻ mice showed partial protection, suggesting that other AhR-expressing antigen-presenting cells—such as macrophages or non-CD11c⁺ DC subsets—may contribute to the immunomodulatory effects of LAQ. These findings support a model in which AhR activation in CD11c⁺ DCs facilitates LAQ-mediated protection, but additional APC populations also participate in this process.

cDC1 cell populations in mice and humans are considered tolerogenic DCs because they express the *Ido1* gene, which drives tryptophan catabolism and generates kynurenines that activate AhR in T cells, promoting regulatory T cell differentiation and suppressing effector T cell responses. Additionally, *Ido2*is expressed in antigen-presenting cells. AhR tightly regulates both genes and is implicated in immune modulation. We further evaluate cDC1 and cDC2 sub-populations (Fig. 4F), observing an increased proportion of cDC1 with a corresponding reduction in cDC2 in CD11c-AhR+ EAU mice treated with LAQ. To address whether this trend resulted from the deletion of AhR or immunization, leading to skewed differentiation of cDC1 and cDC2 populations, we tested naïve CD11c-AhR- and CD11c-AhR+ mice. We did not detect differences in cDC populations among naïve mice (data not shown). This observation supports the idea that LAQ treatment contributes to the differentiation of DCs *in vivo*. Our data support the notion that LAQ treatment may reprogram DC to a tolerogenic phenotype *in vivo* and may thus promote tolerance induction.

### DELAQ inhibits antigen-specific T cell priming and recall responses *in vitro*

Because LAQ retained partial efficacy in CD11c-AhR⁻ mice, we hypothesized that additional AhR-expressing antigen-presenting cell (APC) populations might contribute to its protective effect. To assess the APC-dependent action of DELAQ, we performed in vitro priming assays using irradiated splenocytes as APCs. Naïve CD4⁺ T cells from OTII mice were co-cultured with splenocytes and stimulated with OVA peptide (323–339). DELAQ treatment significantly reduced T cell proliferation, indicating inhibition of antigen-specific T cell priming (Fig. 5A).

**Figure 5.**
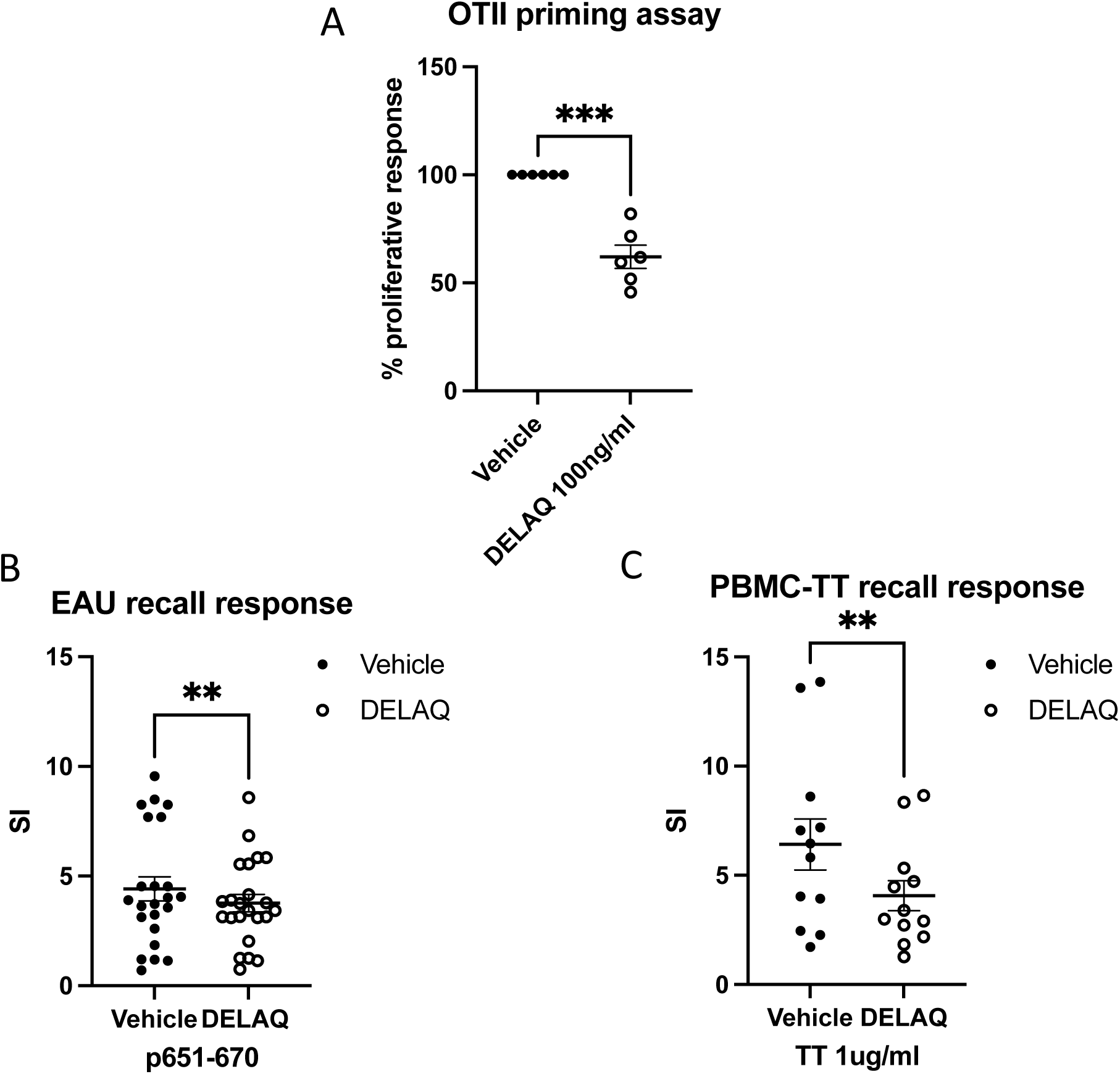
DELAQ inhibits mouse and human antigen-specific T cell responses *in vitro* Antigen-induced proliferative responses of lymphocytes were measured by the incorporation of ^3^H-thymidine during the final 18 hours of incubation, and the stimulation Index (SI) or percentage of proliferative response was calculated. **A -** Naïve CD4+ T cells were purified from OTII mice. Irradiated splenocytes from WT mice were used as antigen-presenting cells. DELAQ at 100 ng/ml or DMSO (vehicle control) was added to the culture with OVA323-339 peptide for 48 h. % of proliferative response = (CPM of cells treated with DELAQ/ CPM of cells with vehicle control) X 100. Combined data from three experiments. **B -** EAU mice were euthanized at the end of the experiments; splenocytes were cultured with IRBP p651-670 for 48 h, in the presence of DELAQ at 100 ng/ml or DMSO only as vehicle control. Each dot represents one mouse. **C -** PBMCs from healthy volunteers were cultured with 1µg/ml of Tetanus Toxoid for four days in the presence of DELAQ at 100 ng/ml or DMSO as vehicle control. Each dot represents one healthy volunteer. t-test was used for statistical comparisons, **p< 0.01 and ***p < 0.001.

To evaluate the effect of DELAQ on previously activated T cells, we next tested its impact on the recall response in EAU mice. DELAQ-treated cultures exhibited reduced proliferation of IRBP-specific T cells harvested at the end of the disease course (Fig. 5B). We extended this analysis to human cells by testing tetanus toxoid–specific recall responses in PBMCs from vaccinated donors. DELAQ again reduced proliferation compared to vehicle controls (Fig. 5C).

Notably, DELAQ had no effect on naïve T cell responses to CD3/CD28 stimulation (Fig. S2B), confirming that its inhibitory effects require the presence of APCs. Together, these findings demonstrate that DELAQ suppresses both priming and reactivation of T cells in an APC-dependent manner, consistent with a mechanism involving AhR-driven modulation of antigen-presenting cells.

## Discussion

This study demonstrates for the first time that Laquinimod (LAQ) is effective in suppressing experimental autoimmune uveitis (EAU). Our data suggest that LAQ interferes with both the priming and effector phases of the autoimmune response, likely through modulation of antigen-presenting cells (APCs), consistent with previous findings in EAE models ^16,17^.

To probe the role of aryl hydrocarbon receptor (AhR) signaling, we examined mice with global or dendritic cell–specific AhR deficiency. AhR-null mice developed full EAU and showed elevated IL-17 and RORγt expression in CD4⁺ T cells, suggesting that AhR is not required for disease initiation but may constrain effector T cell differentiation. These observations prompted in vitro studies using DELAQ, LAQ’s active metabolite. DELAQ activated canonical AhR target genes (*Ahrr*, *Cyp1a1*, *Cyp1b1*) and enhanced Th17 polarization, similar to earlier reports ^17^. In bone marrow–derived dendritic cells (BMDCs), DELAQ induced expression of *Ido1* and *Ido2*, genes associated with tolerogenic APC function.

To examine the cell-type specificity of LAQ, we used CD11c-AhR conditional knockout mice. These mice developed EAU similarly to controls when treated with PBS, suggesting that AhR in dendritic cells is not required for disease induction. However, LAQ conferred strong protection in CD11c-AhR⁺ mice but only partial protection in CD11c-AhR⁻ littermates. These findings implicate dendritic cell–intrinsic AhR signaling in mediating LAQ’s protective effects, though other AhR-expressing APCs likely contribute as well^22^. Consistent with this, LAQ treatment increased the frequency of cDC1 cells in vivo—DCs known for their regulatory potential—and DELAQ induced *Ido1*/*Ido2* expression in BMDCs in vitro.

The ability of LAQ to inhibit EAU even when the disease was induced by adoptively transferred effector T cells was unexpected and led us to test DELAQ’s effects on purified T cells. Using a T cell priming assay with OTII CD4⁺ T cells and irradiated splenocyte APCs, DELAQ significantly reduced antigen-specific proliferation. These results strengthen the conclusion that LAQ interferes with priming, even in preactivated T cells, possibly by modulating recipient APCs in vivo.

To explore translational relevance, we tested DELAQ in human peripheral blood mononuclear cells (PBMCs). As nearly all adults harbor memory T cells specific for tetanus toxoid (TT), this model enabled assessment of recall responses. DELAQ treatment reduced TT-specific proliferation, suggesting that it may dampen effector memory T cell responses in humans. Importantly, DELAQ had no inhibitory effect on purified T cells stimulated through CD3/CD28, confirming that its immunomodulatory activity requires the presence of APCs.

Our in vivo data showed an increase in FoxP3⁺ regulatory T cells (Tregs) in LAQ-treated mice. While we did not directly measure kynurenine levels or demonstrate Treg induction by DCs in vitro, prior studies indicate that IDO-expressing DCs can convert naïve T cells into Tregs through the kynurenine pathway ^30,31^. In our hands, DELAQ did not enhance Treg differentiation in T cell–only cultures, suggesting that APC-mediated signaling is required for this outcome.

Interestingly, LAQ administered from Day 7 post-immunization had limited impact on disease, while treatment of mice receiving activated pathogenic T cells resulted in significant protection. This apparent discrepancy may reflect differing immune contexts. In adoptive transfer, effector T cells may require reactivation in target tissues such as the eye. AhR is expressed in myeloid populations within the eye, and LAQ may disrupt local antigen presentation, similar to effects seen in astrocytes during EAE ^17^. Our in vitro data further support this interpretation: DELAQ suppressed priming and recall responses only when APCs were present.

Although LAQ has previously undergone clinical trials for multiple sclerosis (MS) and Huntington’s disease with only modest or inconclusive efficacy ^25,2^, our findings suggest that autoimmune uveitis may represent a more favorable therapeutic context. Unlike Huntington’s disease, which involves progressive neurodegeneration, uveitis is an inflammatory condition in which disease activity remains dependent on continued immune activation. While MS also involves autoimmune pathology, the disease often includes neurodegenerative components and compartmentalized inflammation in advanced stages ^33,34^, potentially limiting the effectiveness of therapies like LAQ that target early immune events. In contrast, uveitis may allow ongoing recruitment of new effector T cells into the inflammatory pool, particularly during flares, providing a window for LAQ to disrupt priming or reactivation. Moreover, the immune microenvironment of the eye differs from that of the CNS; the blood-retinal barrier may permit more dynamic interaction with immune-modulating therapies ^35^, and ocular antigen presentation may be more accessible to tolerogenic reprogramming. These distinctions, while not guaranteeing therapeutic success, provide a rational basis for considering LAQ in uveitis despite its limited efficacy in other neurological diseases.

In summary, our findings show that LAQ protects against EAU by targeting AhR-expressing APCs to impair both T cell priming and reactivation. DELAQ acts as a biologically active, cell-type–specific AhR ligand that reprograms dendritic cells toward a tolerogenic, IDO-expressing phenotype. These results highlight a mechanism by which AhR signaling can mediate immune tolerance and offer insight into potential therapeutic strategies for human autoimmune disease.

## Acknowledgments

The authors are grateful to Drs. Helena Ericksson and Marie Törngren of Active Biotech AB, Lund, Sweden, for their kind gift of Laquinimod (LAQ) and its derivative, DELAQ, and for a critical review of the manuscript. We thank the NEI Flow Cytometry Core and its staff for their excellent assistance and service.

This study was supported by NIH/NEI Intramural Funding, Project No. EY000184.

---

**Figure S1.**
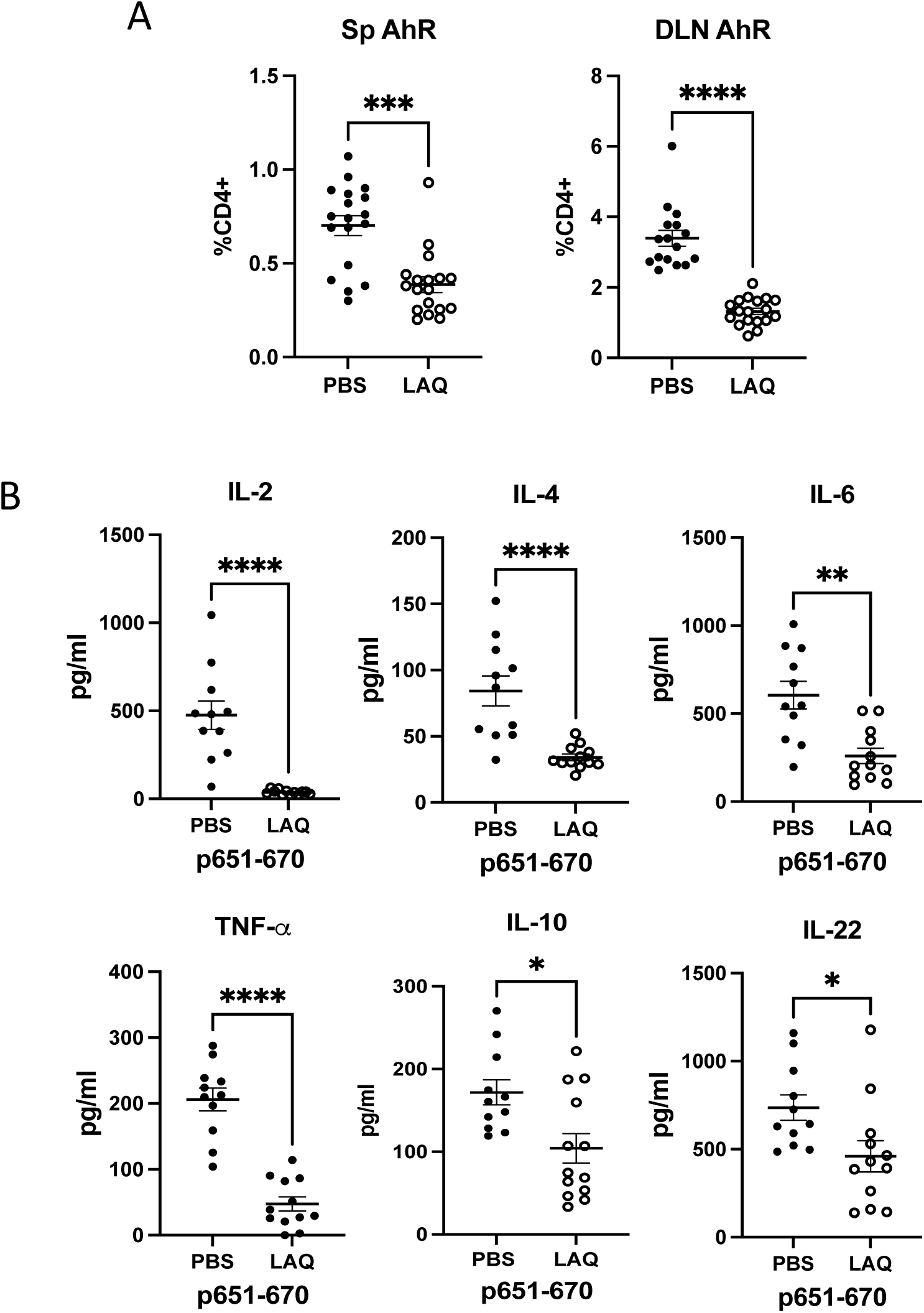
LAQ reduces AhR⁺ cells and broadly suppresses cytokine responses EAU mice were euthanized on day 21 post-immunization. Cells from the spleen (SP) and draining lymph nodes (DLN) were processed for ex vivo testing. **A -** Intracellular AhR expression levels were tested by flow cytometry without stimulation. **B -** IL-2, IL-4, IL-6, TNF-α, IL-10 and IL-22 ELISA on culture supernatants of splenocytes stimulated with p651-670. Each dot represents one mouse, and all data were plotted as mean ± SEM. The Mann-Whitney test was used to determine the significance. *p <0.05, ** p <0.01, *** p <0.001, ****p <0.0001.

**Figure S2:**
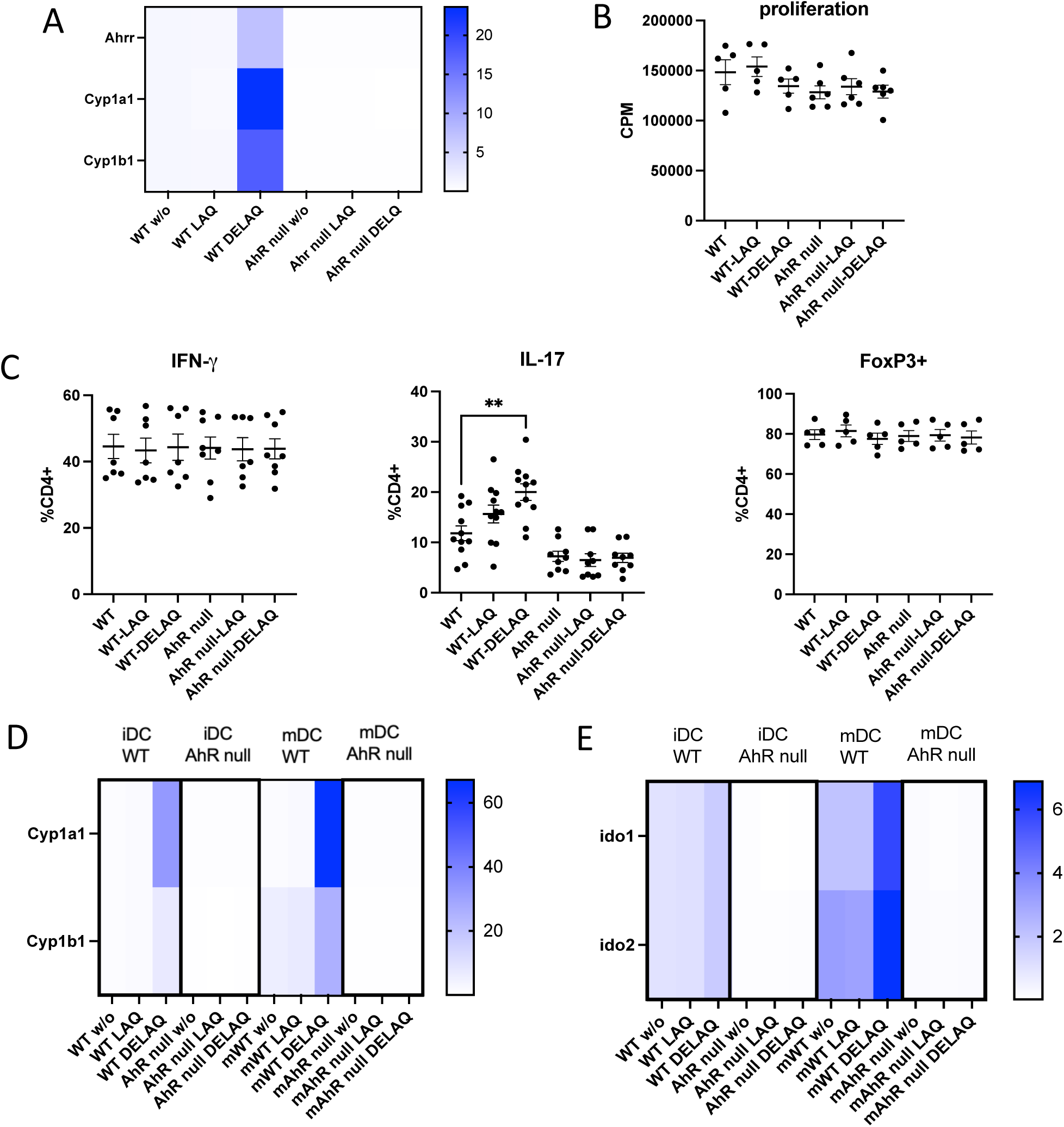
DELAQ activity is AhR-dependent and APC-mediated **A–C**: Naïve CD4+ T cells, purified from WT or AhR null mice, were stimulated with plate-bound anti-CD3 and anti-CD28 antibodies. Cells were treated with LAQ or DELAQ at 100ng/ml for 24 hours. **A –** q-PCR analysis on AhR pathway-related genes, *Ahrr, Cyp1a1,* and *Cyp1b1*. RQ values were plotted. **B -** Proliferative response to LAQ or DELAQ treatment. **C -** cells were polarized to Th1, Th17, and Treg with cytokine cocktails and analyzed by flow cytometry. **D & E**: BMDC generated from B6 and AhR null mice were treated with LAQ or DELAQ at 100 ng/ml for 24 hours. **D -** q-PCR analysis on AhR pathway-related genes *Cyp1a1* and *Cyp1b1*. Combined data from three experiments. **E -** q-PCR analysis on *Ido1* and *Ido2* gene expression. Combined data from three experiments. Each dot represents one experiment. All data were plotted as mean ± SEM. The Mann-Whitney test was used to determine the significance. *p <0.05, **p <0.01, ***p <0.001.

